# Single extracellular vesicle analysis performed by imaging flow cytometry in contrast to NTA rigorously assesses the accuracy of urinary extracellular vesicle preparation techniques

**DOI:** 10.1101/2021.04.01.437817

**Authors:** Marvin Droste, Tobias Tertel, Stefanie Jeruschke, Robin Dittrich, Evangelia Kontopoulou, Bernd Walkenfort, Verena Börger, Peter F. Hoyer, Anja K. Büscher, Basant K. Thakur, Bernd Giebel

## Abstract

Extracellular vesicles (EVs) from several body fluids, including urine, appear as promising biomarkers. Within the last decade, numerous groups have compared the efficacy of EV preparation protocols. Frequently, the efficacy of EV preparation methods is judged by the recovery of particles as estimated by conventional nanoparticle tracking analysis (NTA) or other particle quantification devices. Here, at the example of different urinary EV (uEV) preparation methods, we determined the particle yield in obtained samples with conventional NTA, analyzed their EV content by imaging flow cytometry (IFCM) and quantified the intensity of TSG101 and the contaminant protein uromodulin (UMOD) in Western blots. Our results demonstrate a correlation among CD9-positive objects detected by IFCM and TSG101 Western blot intensities, while particle numbers as determined by NTA correlated with the amount of UMOD.

Consequently, our results question the reliability of conventional NTA analyses for identifying the optimal EV preparation method. Here, in our method comparison, a combination of size exclusion chromatography followed by ultra-filtration showed the highest CD9-positive object and TSG101 protein recovery, and in relation to the number of CD9-positive objects, the lowest amount of UMOD contamination.

## Introduction

Small extracellular vesicles (sEVs) such as exosomes (70-150 nm) are membrane-coated particles containing nucleic acids, proteins and lipids of cellular origin (1). Due to their cell type specific assembly, they have been qualified as biomarkers for various diseases (2). sEVs can be detected in all biological fluids, including urine (3). As the collection of urine is non-invasive and enables acquisition of large sample volumes, small urinary EVs (uEVs) provide an easily accessible source for the identification of novel biomarkers (4-6). However, the applied uEV preparation technique largely influences the yield and purity of uEVs and thus the validity of associated biomarkers. Although the *International Society for Extracellular Vesicles* (ISEV) urges researchers to proceed towards more standardized EV isolation protocols (7), the variety of techniques applied is large. Differential centrifugation procedures including ultracentrifugation (UC) were the gold standard for sEV preparation for years (8). However, as it has been recognized that UC can promote EV aggregation, does not achieve sufficient purity and may result in insufficient EV enrichment (9, 10), alternative methods are increasingly used for sEV preparation, including polymer-based precipitation, size exclusion chromatography (SEC) and filtration-based methods (11, 12). Concerning the preparation of uEVs, various challenges need to be considered. In addition to the varying concentration of void urine and interindividual differences in uEV profiles (13), especially the presence of highly abundant proteins, such as uromodulin (Tamm Horsfall protein, UMOD), hampers the purification of uEVs (14-17). Although several groups already compared different uEVs enrichment methods and have also focused on the reduction of co-prepared proteins, no consensus has been reached to date (18-23).

Critical parameters in evaluating the accuracy of EV preparation methods depend on the analysis technologies being used for the characterization of obtained EV samples. Currently, the validity of EV preparation methods is frequently judged by particle quantification technologies, e.g., by nanoparticle tracking analysis (NTA), which we and others introduced in 2011 as an “exosome” quantification method (24, 25). However, upon comparing NTA with imaging flow cytometry (IFCM) analysis, the latter allowing single EV analyses even in non-processed EV containing samples (26-28), we had to learn that depending on the initial sample material and the applied preparation method, obtained EV samples contain far more particles than EVs. Indeed, protein aggregates that are formed in urine appear in NTA as small particles which are indistinguishable from uEVs (29). Consequently, it needs to be considered that results from NTA and other particle quantification devices in their original form can be misleading, especially if the accuracy and efficacy of EV preparation methods is compared.

Being interested in qualifying EVs as novel urine-derived biomarkers, we thus decided to re-assess different uEV preparation methods. Now, in addition to the classical analysis techniques, i.e., NTA, Western blot (WB) and transmission electron microscopy (TEM), we performed IFCM analyses for the detection and semi-quantification of single uEVs. To this end, we at first searched for appropriate antibodies allowing us to identify a prominent proportion of uEVs in fresh void urine. For practical reasons, it is often required to work with stored void urine samples. Consequently, we also studied the impact of different storage temperatures on the recovery of antibody-labelled uEVs. Within the method comparison, we focused on the recovery and purity of uEVs which were successfully labelled with the selected antibody. According to the results of the IFCM analyses, other uEV preparation techniques appeared favorable than those which would have been selected according to the results of the NTA analyses. Since obtained IFCM data are supported by the results of the WB and TEM, but to a lesser extent by the NTA data, we like to conclude that results of the IFCM analyses are more accurate for the characterization of EV samples and for the evaluation of EV preparation protocols than those obtained with NTA or other conventional particle quantification devices.

## Material and Methods

### Urine sample collection and general preparation

Void urine samples were collected from healthy adult male and female volunteers following obtained written informed consent. The study was approved by the ethics committee of the Medical Faculty of the University of Duisburg-Essen (no. 18-8494-BU). After collection, samples were centrifuged at 500 × *g* for 10 min (4°C) and 3,000 × *g* for 20 min (4°C) to remove cells, cellular debris and larger vesicles. Supernatants were stored at −80°C until further processing unless otherwise indicated. Details of the centrifugation procedures are provided in Supplementary Table S4. Urine samples considered for IFCM analyses were pre-cleared by filtration using 0.22 µm polyethersulfone syringe filters (Minisart, Sartorius, Göttingen, Germany).

### Polyethylene glycol precipitation followed by ultracentrifugation (PEG-UC)

PEG precipitation and subsequent UC was performed as described previously (30) with some minor modifications. Briefly, pre-processed urine samples were spun for 45 min at 10,000 × *g* and 4°C. Following addition of 1 mL 50 w/v % PEG-6000 (Sigma-Aldrich, Steinheim, Germany) and 750 µL of 0.9% NaCl (Fresenius Kabi, Bad Homburg, Germany) to 8.25 mL supernatant (final PEG concentration: 10%), uEVs were precipitated at 4°C overnight. Precipitates were pelleted by centrifugation for 30 minutes at 1,500 × *g* and 4°C. Pellets were resuspended in 0.9% NaCl and spun at 110,000 × *g* for 2 hours at 4°C in an XPN-80 ultracentrifuge equipped with a Type 50.4 Ti rotor (Beckman Coulter, Krefeld, Germany; k-factor: 93). UC pellets were resuspended in 1 mL of 0.9% NaCl and stored at −80°C until further analysis.

### PEG precipitation followed by SEC (PEG-SEC)

Exactly as before, urine samples were pre-processed. 200 µL of PBS (ThermoFisher Scientific / Gibco, Carlsbad, CA, USA) were added to 7.8 mL pre-processed urine. Following addition of 2 mL 50% w/v PEG-6000 solution (Sigma-Aldrich; final concentration: 20%) uEVs were precipitated at 4°C overnight. Precipitates were pelleted by centrifugation for 35 minutes at 1,500 × *g* and 4°C. Next, size exclusion chromatography was performed with 10 mL of Sepharose CL-2B (GE Healthcare, Uppsala, Sweden) using self-packed and PBS-equilibrated columns (Econo-Pac 20 mL, Bio-Rad Laboratories, Hercules, CA, USA) according to the “Mini-SEC” procedure reported by Hong and colleagues (31). The resuspended uEV pellet was applied onto the top mesh of the size exclusion columns. 6 fractions, each of 1 mL, were eluted. After elution of each fraction, 1 mL PBS was added to the column. Fraction 4, the fraction containing most EVs, was used for all downstream analyses.

### UC followed by size exclusion chromatography (UC-SEC)

For the UC-SEC-method, pre-processed urine samples were spun for 45 minutes at 17,000 × *g* at 4°C. Next, supernatants were spun for 70 minutes at 200,000 × *g* and 4°C in an XPN-80 ultracentrifuge equipped with a Type 50.4 Ti rotor (Beckman Coulter, Krefeld, Germany; k-factor: 51). Obtained pellets were resuspended and washed in 2 mL PBS, followed by pelleting uEVs again by repeating the UC step. SEC of resuspended pellets was performed exactly as described before.

### Ultrafiltration followed by SEC (UF-SEC)

UF-SEC was performed according to the protocol by Monguió-Tortajada et al. (32) with some minor modifications. 10 mL of pre-processed urine was spun at 17,000 × *g* for 15 minutes at 4°C. The supernatant was then supplemented with 5 mL of 0.9% NaCl and concentrated by centrifugation (4,000 × *g*; 10 min; room temperature) using Amicon Ultra-15 100 kDa centrifugal filtration units (regenerated cellulose; Merck/Millipore, Cork, Ireland). To further reduce the protein content, the concentrate above the filter was diluted with 0.9% NaCl to a final volume of 10 mL and re-concentrated by centrifugation. Concentrates were harvested and supplemented with 0.9% NaCl to a final volume of 1 mL. Thereafter, SEC was performed applying the same protocol than for UC-SEC and PEG-SEC.

### Ultrafiltration followed by immunoaffinity capturing (UF-MACS)

Immunoaffinity isolation of uEVs was performed using a commercially available kit (Exosome Isolation Kit Pan, human, Miltenyi Biotec, Bergisch Gladbach, Germany) according to the manufacturer’s instruction. The kit is based on the immunomagnetic isolation of uEVs carrying any of the surface epitopes CD9, CD63 or CD81. Briefly, pre-processed void urine was pre-cleared by centrifugation at 10.000 × *g* for 30 minutes (4°C). Then, 10 mL of the supernatant were concentrated using an Amicon Ultra-15 100 kDa filter (regenerated cellulose; Merck/Millipore) by centrifuging at 4.000 × *g* for 10 minutes. The concentrate was adjusted to 2 mL with 0.9% NaCl. Then, 50 µl of antibody-loaded magnetic beads were added. After incubation for 1 hour at room temperature the sample was loaded onto an equilibrated separation column within a magnetic stand (MACS MultiStand with µMACS separator, Miltenyi Biotec). After serial washing, uEV-bead-aggregates, the column was removed from the magnetic stand. uEV-bead aggregates were eluted by flushing with 100 µl isolation buffer. The samples were adjusted to 1 mL with 0.9% NaCl.

### Membrane affinity-based isolation (ExoEasy)

The commercial exoEasy Maxi Kit (Qiagen, Hilden, Germany) was used for membrane affinity-based isolation of uEVs according to the instructions of the manufacturer. Briefly, pre-cleared void urine was spun at 10.000 × *g* for 45 minutes (4°C). The supernatant was mixed 1:1 with the provided binding buffer and applied to the provided centrifugation columns. After centrifugation, membranes were washed, and bound components were eluted by the addition of 1 mL of the provided elution buffer and subsequent centrifugation.

### Imaging Flow Cytometry (IFCM)

IFCM was performed on the AMNIS ImageStreamX Mark II Flow Cytometer (AMNIS/Luminex, Seattle, WA, USA) as described before (26, 28). Details for all antibodies used are provided in Suppl. Table S5. Generally, antibody incubation time was 1 hour at room temperature. According to the recommendations of the MIFlowCyt-EV guidelines (33), unstained uEV samples, NaCl-HEPES buffer with antibodies but without uEV sample as well as stained sample supplemented with 1% NP40 (Calbiochem, San Diego, CA, USA) were analyzed as controls. After staining, samples were diluted with PBS and analyzed using the built-in autosampler for 96-well round bottom plates. Acquisition time was selected as 5 minutes per well. Data were acquired at 60x magnification, low flow rate and with removed beads option deactivated. Further details are provided in Suppl. Tables S6-7.

Data were analyzed as described previously using the IDEAS software (version 6.2) (26, 28). Fluorescent events were plotted against the side scatter (SSC). A combined mask feature was used (MC and NMC) to improve the detection of fluorescent images. Images were analyzed for coincidences (swarm detection) by using the spot counting feature. Every data point with multiple objects was excluded from the analyses. Events with low side scatter values (< 500) and fluorescence intensities higher than 300 were considered as uEVs. Average concentrations were calculated according to the acquisition volume and time.

### Nanoparticle tracking analysis (NTA)

Average size distribution and particle concentration analyses of uEV samples were performed by NTA on the ZetaView PMX-120 platform equipped with the software version 8.03.08.02 (ParticleMetrix, Meerbusch, Germany) exactly as described before (30). Briefly, samples were diluted in NaCl. 1 mL diluted sample volume was loaded into the flow cell and recorded for 55 seconds. Particle sizes and numbers of all 11 positions were recorded and calculated as the mean of the results. The following settings were used: positions: 11; cycles: 5; quality: medium; min. brightness: 20; min. size: 5; max. size: 200; trace length: 15; sensitivity: 75; shutter: 75; framerate: 30.

### SDS-PAGE / Immunoblot

Equal volumes of each sample were lysed in radioimmunoprecipitation assay buffer (1:1; 150 mM NaCl, 50 mM Tris, 1% Triton X-100, 0.1% SDS, 0.5% sodium desoxycholate) containing protease inhibitors (1:100; Pefabloc SC, Sigma-Aldrich). Proteins were separated under reducing and non-reducing conditions on Mini-Protean TGX Any-kD precast gels (Bio-Rad, Hercules, CA, USA) and transferred to nitrocellulose membranes (GE / Amersham, Buckinghamshire, UK) using a Fastblot B34 blotting device (Biometra, Göttingen, Germany). Membranes were blocked with 5% milk powder (Sigma-Aldrich) solved in PBS-T. Membranes were incubated with primary antibodies (Suppl. Table S5) overnight at 4°C. Following serial washing, bound antibodies were counterstained with horseradish peroxidase-conjugated secondary antibodies (Suppl. Table S5) for 1 hour at room temperature. After washing, the Super Signal West Femto Chemiluminescent Substrate (Thermo Scientific, Rockford, IL, USA) was applied according the manufacturer’s instructions. Obtained signals were documented with the Fusion FX7 detection system (Vilber Lourmat, Eberhardzell, Germany).

### Transmission Electron Microscopy (TEM)

A carbon coated formvar film supported by a 200-mesh copper grid (Plano, Wetzlar, Germany) was pretreated with glow discharging agent (Ted Pella, Redding, CA, USA) to create a hydrophilic surface. 5 µl of each given sample suspension was placed on a grid and incubated for 5 minutes. The grids were washed 3 times for 1 minute by placing it on 30 µl-droplets of deionized water, before they were stained for 1 minute on a 20 µl droplet of 1.5% aqueous phosphotungstic acid solution (w/v, Carl Roth, Karlsruhe, Germany). Thereafter, following removing excess staining solution with a piece of filter paper, they were dried at ambient air. Images were acquired using a JEOL JEM 1400Plus (JEOL, Tokyo, Japan), operating at 120 kV and equipped with a 4096×4096 px CMOS camera (TVIPS, Gauting, Germany). The image acquisition software EMMENU (Version 4.09.83) was used for taking 16-bit images. ImageJ software (Version 1.52b, https://imagej.nih.gov/ij/) was used to process and analyze obtained images.

### Data Analysis and Statistics

Computational data plotting, analysis and visualization was performed using Microsoft Excel 2019 and GraphPad Prism (version 8.4.0). Image analysis, including band densitometry, was performed using ImageJ. Band density was calculated by measuring the density of a region of interest (ROI) placed around the bands after subtraction of background density. Statistical significance was calculated using unpaired Student’s t-test and, in case of more than two groups, a one-way ANOVA with Tukey’s post-test was applied. Data correlation analysis was performed and plotted using Pearson’s correlation coefficient with two-tailed Student’s t-test. Correlation analyses for particle numbers and CD9-positive objects was performed in RStudio (version 1.4.1103). P values < 0.05 were considered as significant. Measurement results are given as means ± standard deviation, unless otherwise indicated.

### Data Reporting

We have submitted all relevant data of our experiments to the EV-TRACK knowledgebase (EV-TRACK ID: EV210067) (34).

## Results

### CD9 is abundantly present on small urinary EVs of healthy donors

Recently, we set up protocols to label sEVs with fluorescent conjugated anti-CD9, anti-CD63 and anti-CD81 antibodies in otherwise non-processed cell culture supernatants, allowing us to immediately analyze the labeled EVs by IFCM (26, 28). Being interested in uEVs in the context of biomarker research, we explored whether a comparable labeling technology could be used for the detection of uEVs in fresh void urine samples of healthy donors (Fig. 1). Applying the established protocols, we recovered discrete CD9^+^ uEV populations in all urine samples tested (n = 4). In contrast, CD63^+^ and CD81^+^ uEV populations were hardly detectable (CD9: 4.1×10^5^ ± 4.8×10^5^ objects/mL; CD63: 1.8×10^4^ ± 1.4×10^4^ objects/mL; CD81: 6.8×10^4^ ± 1.0×10^5^ objects/mL). Thus, CD9, but neither CD63 nor CD81, presents an abundant uEV marker in fresh human void urine.

**Fig. 1:**
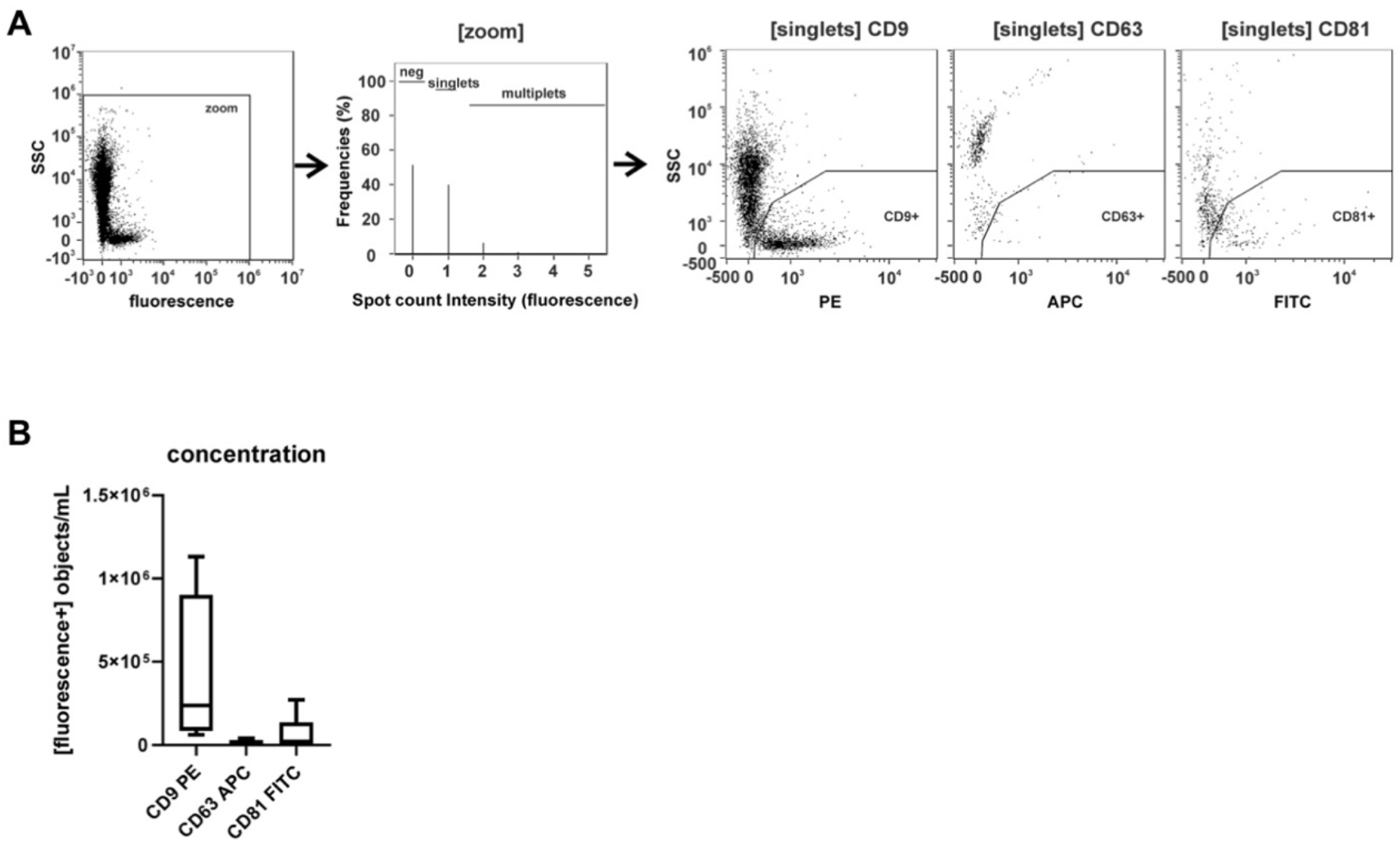
CD9 is abundantly present on small urinary EVs of healthy donors. Freshly pre-processed void urine samples (n = 4) were stained with anti-CD9, anti-CD63 and anti-CD81 antibodies and after 1h immediately analyzed by imaging flow cytometry (IFCM). **A** Applied gating strategy: of all recorded signals (1^st^ plot), data points lacking any spot count signal or showing coincidences were neglected (2^nd^ plot). Side scatter intensities of single objects are plotted against the fluorescence intensities of CD9-, CD63- and CD81-labeled objects. **B** Box plots reflecting the numbers of recorded CD9^+^, CD63^+^ and CD81^+^ objects of the different experiments.

### Storage temperature affects the recovery of urinary sEVs

Commonly, biomarker screening projects depend on preserved donor samples. Regularly, urine samples of healthy donors and patients are cryopreserved either at −20°C or at −80°C. To test whether cryopreservation affects the quality of respective samples, we evaluated the impact of the storage temperature on the recovery of CD9^+^ uEVs (Fig. 2). To this end, 5 fresh void urine samples of healthy donors were processed by low-speed centrifugation and either stored for 1 month at room temperature, 4°C, −20°C or −80°C, or analyzed immediately. For uEV marker analyses, stored as well as fresh samples were filtered through 0.22 µm polyethersulfone membrane filters and stained with anti-CD9 antibodies. Stained samples were analyzed by IFCM. Notably, compared to the freshly prepared urine sample, the number of CD9^+^ uEVs declined under all storage conditions, with the lowest decline in samples that had been stored at −80°C (mean recovery: 36.5% ± 8.0%; mean decline vs. no storage: 4.3×10^5^ CD9^+^ objects/mL; p = 0.0001***) (Fig. 2). The highest decline was observed in samples stored at −20°C (mean recovery: 4.8% ± 2.9%); mean decline vs. no storage: 6.5×10^5^ CD9^+^ objects/mL; p < 0.0001**** (Fig. 2). The difference between the mean recoveries of uEVs stored at −20°C vs. −80°C was statistically significant (p < 0.0001****) (Fig. 2). Thus, if urine samples cannot be immediately processed for uEV analyses, our results indicate that they are ideally stored at −80°C.

**Fig. 2:**
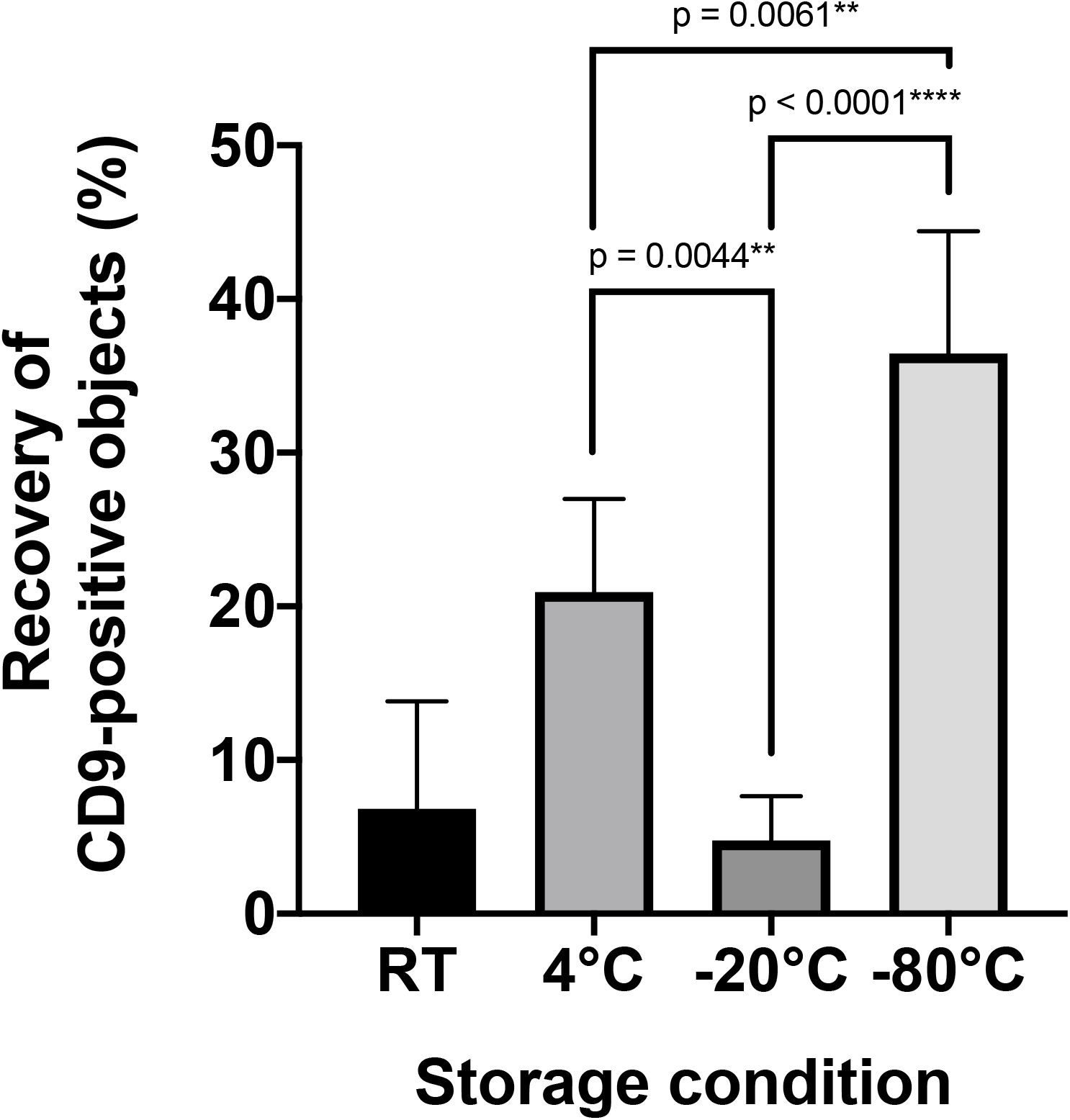
Recoveries rates of CD9^+^ uEVs in void urine samples depend on the storage temperature. Freshly prepared, cell-free void urine samples (n = 5) were analyzed by IFCM immediately after antibody staining with anti-CD9 antibodies (no storage) or after storage for one month either at room temperature, (RT, 20°C), 4°C, −20°C, or −80°C. The recovery of CD9^+^ objects in stored samples was calculated as the ratio of CD9^+^ objects before and after storage.

### Imaging flow cytometry and NTA analyses provide incongruent results regarding the efficacy of uEV preparation methods

To evaluate the accuracy of different uEV preparation methods we used urine samples that had been stored at −80°C. Five independent void urine samples, two more aqueous and three more concentrated ones, were processed, each with five different protocols frequently applied to enrich uEVs: i) polyethylene glycol-precipitation followed by ultracentrifugation (PEG-UC) (30) or ii) size exclusion chromatography (PEG-SEC) (19); iii) UC followed by SEC (UC-SEC) (15); iv) ultrafiltration followed by SEC (UF-SEC) (32); and v) by a one-step protocol using the commercial ExoEasy Maxi Kit based on membrane affinity (Fig. 3). The first void urine sample was also processed with a sixth method, ultrafiltration followed by immunoaffinity capturing (UF-MACS) (35). Due to the presence of EV-microbead aggregates, this sample could neither be analyzed by IFCM nor by NTA. Consequently, MACS preparations were not performed for the remaining four void urine samples.

**Fig. 3:**
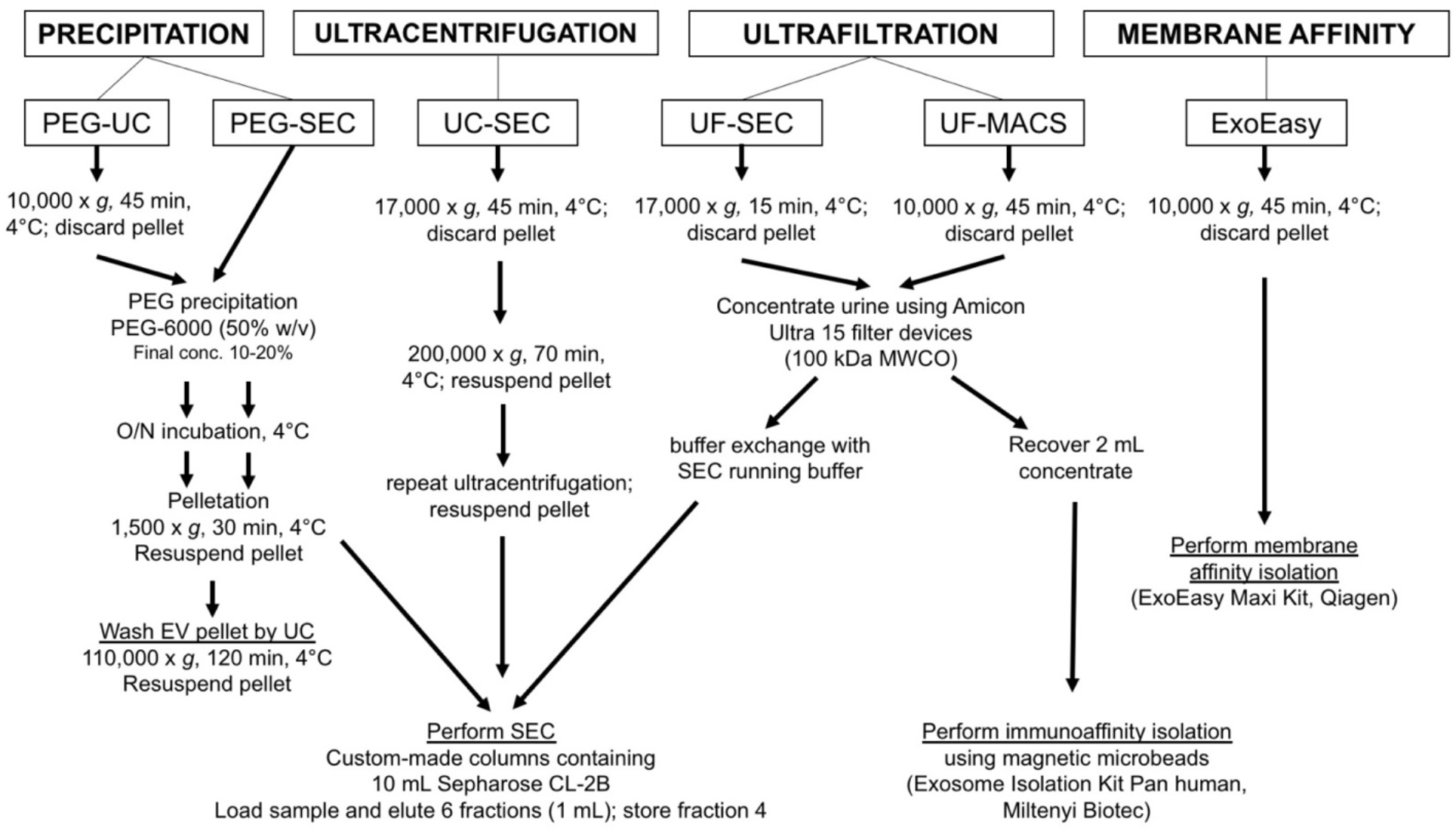
Experimental design of the method comparison using 5 different methods for uEV preparation of 3 concentrated and 2 aqueous void urine samples and a sixth method for one of the normal void urine samples. According to published protocols, void urine samples were differently pre-processed, either at 10,000 × *g* or at 17,000 × *g*. All details for the next procedures are provided. Following uEV preparation, uEV samples were analyzed by IFCM, NTA, Western blot and transmission electron microscopy. Methods are PEG precipitation followed by ultracentrifugation (PEG-UC): (30), PEG precipitation followed by size exclusion chromatography (PEG-SEC) (19), ultracentrifugation followed by size exclusion chromatography (UC-SEC) (15), ultrafiltration followed by size exclusion chromatography UF-SEC (32) and ExoEasy Maxi Kit, Qiagen, based on membrane affinity (ExoEasy). The sixth method performed on one of the normal void urine samples was ultrafiltration followed by immunoaffinity capturing with commercial anti-CD9, anti-CD63 and anti-CD81 antibody-loaded magnetic beads (UF-MACS) (35).

Following uEV enrichment, samples were at first analyzed for their CD9^+^ uEV content by IFCM and then for total particles by NTA. IFCM analyses of the 5 different samples with the five preparation protocols revealed that the highest CD9^+^ object numbers were obtained with the UF-SEC method (2.5×10^6^ ± 2.2×10^6^ objects/mL), followed by UC-SEC (4.0×10^5^ ± 2.7×10^5^ objects/mL). The number of the detected CD9^+^ objects of PEG-SEC, PEG-UC and ExoEasy preparations were close to the IFCM detection limit (PEG-SEC: 8.2×10^4^ ± 7.6×10^4^ objects/mL; PEG-UC: 7.9×10^4^ ± 3.1×10^4^ objects/mL; ExoEasy: 3.0×10^4^ ± 2.0×10^3^ objects/mL) (Fig 4A, Suppl. Fig. S1, S2A). Upon focusing on potential differences between more concentrated and aqueous samples, UC-SEC and UF-SEC yielded similar numbers of CD9^+^ objects from the more aqueous void urine samples, while the CD9^+^ object yield of concentrated void urine samples was much higher following UF-SEC than following UC-SEC (Fig. 4B).

**Fig. 4:**
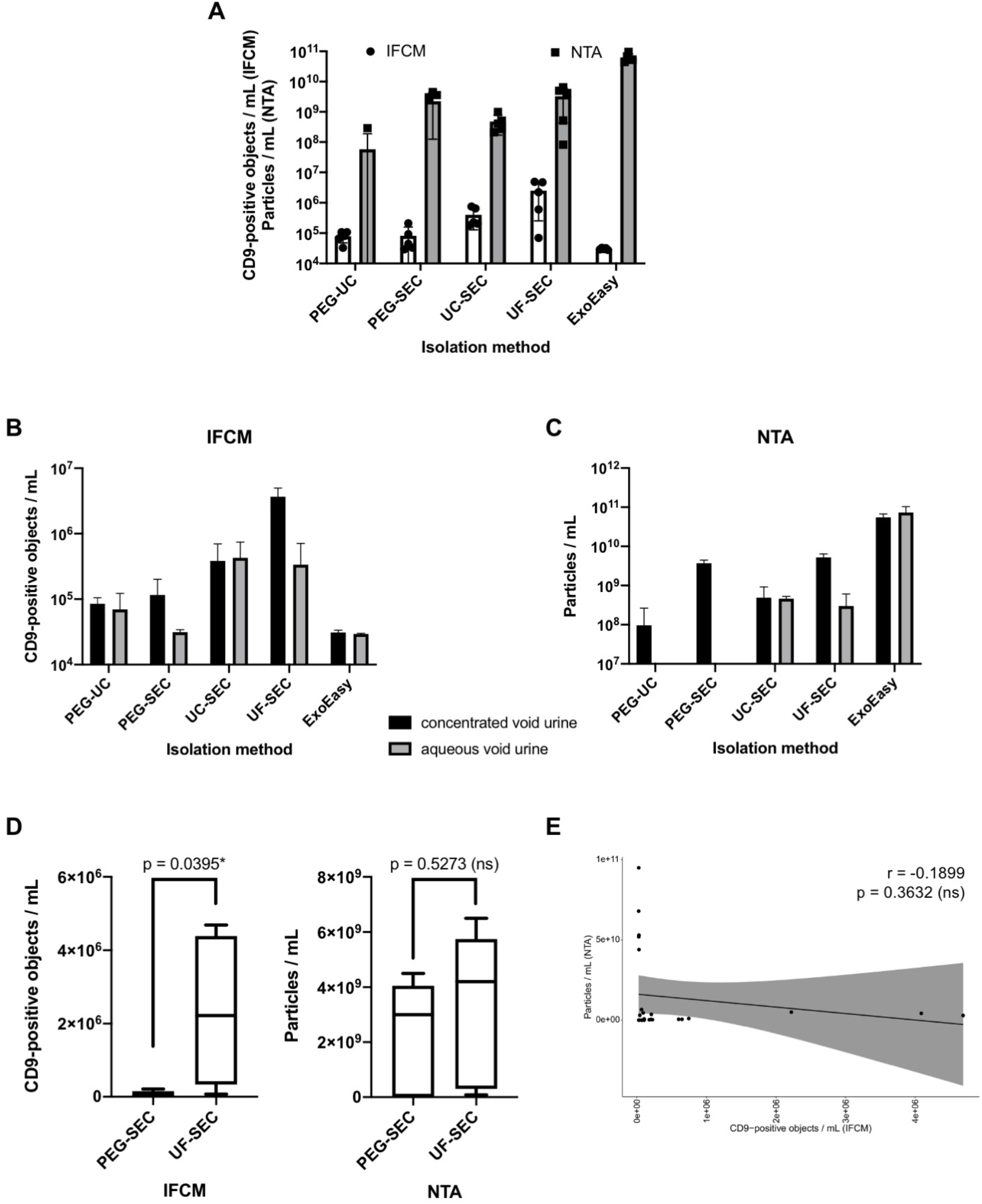
CD9^+^ object recovery rates determined by IFCM are incongruent to the particle recovery rates determined by NTA. Following processing of 5 void urine samples with the methods shown in Fig. 3 (3 concentrated and 2 aqueous void urines), obtained samples were analyzed for the presence of CD9^+^ objects by IFCM and for the presence of particles by NTA. **A** Side-by-side comparison of average numbers of CD9^+^ objects detected by IFCM and average particle numbers as quantified by NTA. **B** Comparison of average numbers of CD9^+^ objects detected by IFCM in uEV samples prepared with the 5 different methods from normal (n = 3) and aqueous (n = 2) void urine, **C** Comparison of average particle numbers detected by NTA in uEV samples prepared with the 5 different methods from normal (n = 3) and aqueous (n = 2) void urine. **D** Side by side comparison of average CD9^+^ object numbers (left) and average particle numbers (right) detected in UF-SEC and PEG-SEC samples (same data as shown in A). Statistical analysis was performed with the Student’s t-test. **E** Pearson’s correlation analysis of all recorded IFCM and NTA data (n = 25).

All samples were also analyzed by NTA. The highest particle numbers were recorded in EV samples prepared with the ExoEasy Kit (6.2×10^10^ ± 2.0×10^10^ particles/mL), followed by those prepared with UF-SEC (3.3×10^9^ ± 2.8×10^9^ particles/mL) and PEG-SEC (2.2×10^9^ ± 2.1×10^9^ particles/mL). Slightly lower particle concentrations were recorded in UC-SEC (4.8×10^8^ ± 3.1×10^8^ particles/mL) and PEG-UC (5.8×10^7^ ± 1.3×10^8^ particles/mL) samples (Fig. 4A). Notably, high particle numbers were obtained with the PEG-SEC method from concentrated but not from aqueous void urine samples (Fig. 4C, Suppl. Fig. S2B).

Thus, although both methods are intended to analyze single EVs, the results are incongruent. The results of IFCM identified the UF-SEC method as the method allowing the highest CD9 yield, while according to NTA analysis the highest particle yield was obtained with the ExoEasy protocol. We observed that in general all samples contained much more particles being measured by NTA than CD9^+^ objects measured by IFCM (Fig. 4). The difference is highlighted upon comparing the NTA and IFCM data of the UF-SEC and PEG-SEC samples. While according to the NTA data no significant difference was documented (p = 0.5273, Fig. 4D), IFCM detected significantly lower CD9^+^ objects in PEG-SEC than in UF-SEC samples (p = 0.0395*). Notably, IFCM and NTA data of all analyzed samples showed no significant correlation (r = −0.1899, p = 0.3632), demonstrating that the results obtained with the two analysis methods are not congruent (Fig. 4E).

### Western blot results correlate with results from Imaging Flow Cytometry but not with those of NTA

To comply with the MISEV2018 criteria (7) and to potentially solve the inconsistency among the IFCM and NTA data, Western blot (WB) analyses were performed, initially with freshly prepared EV samples obtained from the first void urine sample investigated, one of the more concentrated void urines. These EV preparations, including the aforementioned UF-MACS sample, were separated under reducing and non-reducing conditions. The membrane of the reducing WB was sequentially probed with anti-uromodulin (UMOD) and anti-TSG101 antibodies and that of the non-reducing WB with anti-CD9 antibodies (Fig. 5A). UMOD, an abundant contaminating urinary protein, was detected in high amounts in PEG-SEC and ExoEasy samples and lower amounts in UC-SEC and UF-SEC samples. Hardly any UMOD was recovered in the UF-MACS sample. Unexpectedly, no bands, neither for UMOD nor for TSG101 and CD9, were detected in the PEG-UC preparation. By far the highest TSG101 content was obtained by the UF-SEC method, followed by ExoEasy, UF-MACS and PEG-SEC. Hardly any TSG101 was recovered applying UC-SEC. UF-SEC and UF-MACS contained comparable CD9 contents, which were higher than in the PEG-SEC sample. No CD9 was recovered in UC-SEC and ExoEasy samples (Fig. 5A).

**Fig. 5.**
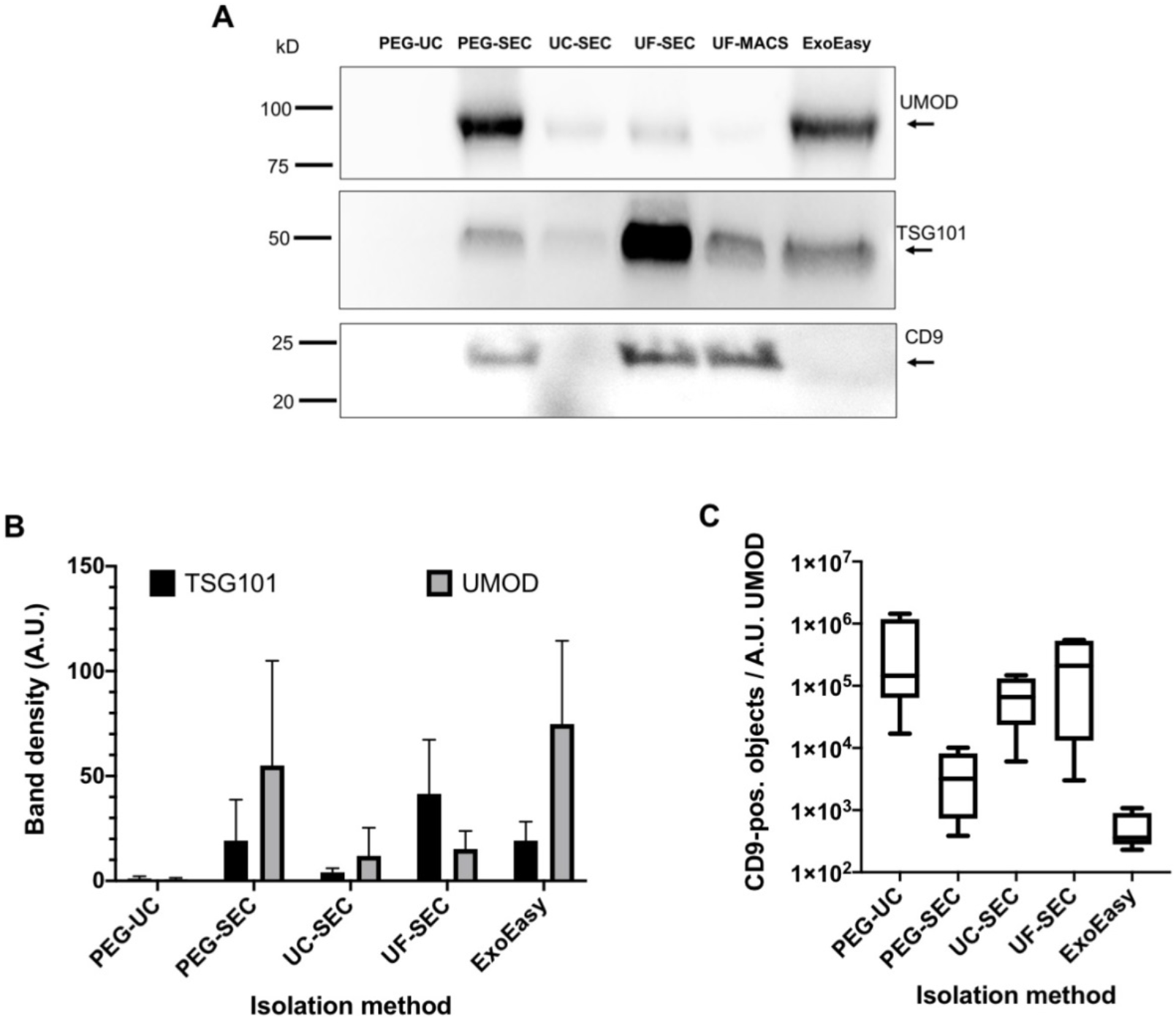
The UF-SEC uEV preparation method outcompetes the other uEV preparation methods in terms of EV marker protein recovery and purity. Western blot results of one of the processed void urine samples performed on freshly obtained uEV samples. Sample loading was adjusted to volume equivalents of the initial void urine sample. uEV samples were separated under reducing and non-reducing conditions. The Western blot from the reducing condition was probed with anti-TSG101 and anti-UMOD antibodies, the Western blot from the non-reducing conditions was probed with anti-CD9 antibodies. Bands were visualized after counterstaining with HRP-conjugated secondary antibodies and addition of chemiluminescent HRP substrate. **B** Comparison of average TSG101 and UMOD band intensities of all uEV samples arranged according to the applied purification method. All samples were stored at −80°C before Western blot analysis, images of all relevant Western blots are shown in Suppl. Fig. S3. **C** Average sample purities calculated as the ratio of the average TSG101 to the average UMOD band intensities.

To substantiate the data, reducing and non-reducing WBs were performed of all obtained EV samples in parallel, including the samples which had initially been tested in WB. In none of the samples, all of which had been stored at −80°C before, CD9 was detected in WB, not even in the uEV samples of the first void urine which had shown CD9 bands in the initial WB when the EV samples were freshly prepared. Since CD9 was well detected in these WBs in the control lanes, the results of the WBs are trustable, supporting unpublished discussions in the field that storage of uEV samples can affect the detectability of certain marker proteins including CD9.

In contrast, the reducing WBs showed clear TSG101 und UMOD bands. The band intensities of the uEV samples of the first processed void urine were comparable in the WB before and after uEV storage. Consequently, we focused on TSG101 and UMOD WB data in subsequent analyses (Suppl. Fig. S3).

The intensities of the TSG101 bands were quantified by densitometry (Fig. 5B). Notably, the WBs of the more concentrated void urine samples showed comparable results to the initially processed concentrated void urine sample, thus favoring UF-SEC as the method of choice for the highest uEV recovery. However, consistent to the results of the IFCM analyses, results of the more aqueous void urine samples were different. Resulting TSG101 bands were weaker than in the samples obtained from concentrated urine samples. Neither TSG101 nor UMOD was recovered in PEG-prepared samples, irrespective of whether they were subsequently processed by SEC or UC (HD4, HD5, Suppl. Fig. 3). In contrast to the samples prepared from more concentrated void urine samples, UF-SEC and ExoEasy samples now contained comparable amounts of TSG101 that were in the same range as in the ExoEasy samples prepared from the concentrated void urine samples. Notably, from the aqueous void urine samples, TSG101 was also recovered using UC-SEC, albeit in lower concentrations than in the UF-SEC and ExoEasy samples. As in preparations from concentrated void urine samples, the UMOD concentration was lower in UF-SEC samples than in the ExoEasy samples. Thus, as in the uEV samples obtained from the concentrate void urines, the ratio between the intensity of the TSG101 to the UMOD band was highest in UF-SEC preparations (due to the absence of TSG101 bands, no such ratios were calculated for PEG processed samples) (Fig. 5C). Overall, our results demonstrate that for more aqueous as well as concentrated void urine samples UF-SEC is the best method tested here.

Next, we correlated the protein concentration of TSG101 and UMOD derived from all densitometric WB analyses with corresponding IFCM and NTA data. While a significant correlation among TSG101 band density and the numbers of CD9^+^ objects measured by IFCM were found (r = 0.7531, p < 0.0001****; Fig. 6A), the TSG101 band densities did not correlate with the numbers of particles measured by NTA (r = 0.1289, p = 0.5392; Fig. 6B). Remarkably, a correlation among the intensities of UMOD bands and particle numbers was obtained (r = 0.6314, p = 0.0007***; Fig. 6C), while no correlation was recognized among the UMOD concentration and the number of CD9^+^ objects measured by IFCM (r = -0.2457, p = 0.06038; Fig. 6D). Thus, our data indicate that NTA numbers are significantly affected by impurities, while impurities apparently do not affect results of IFCM. Consequently, IFCM analyses assess EV contents in samples prepared with different methods much more reliable than conventional NTA analyses.

**Fig. 6.**
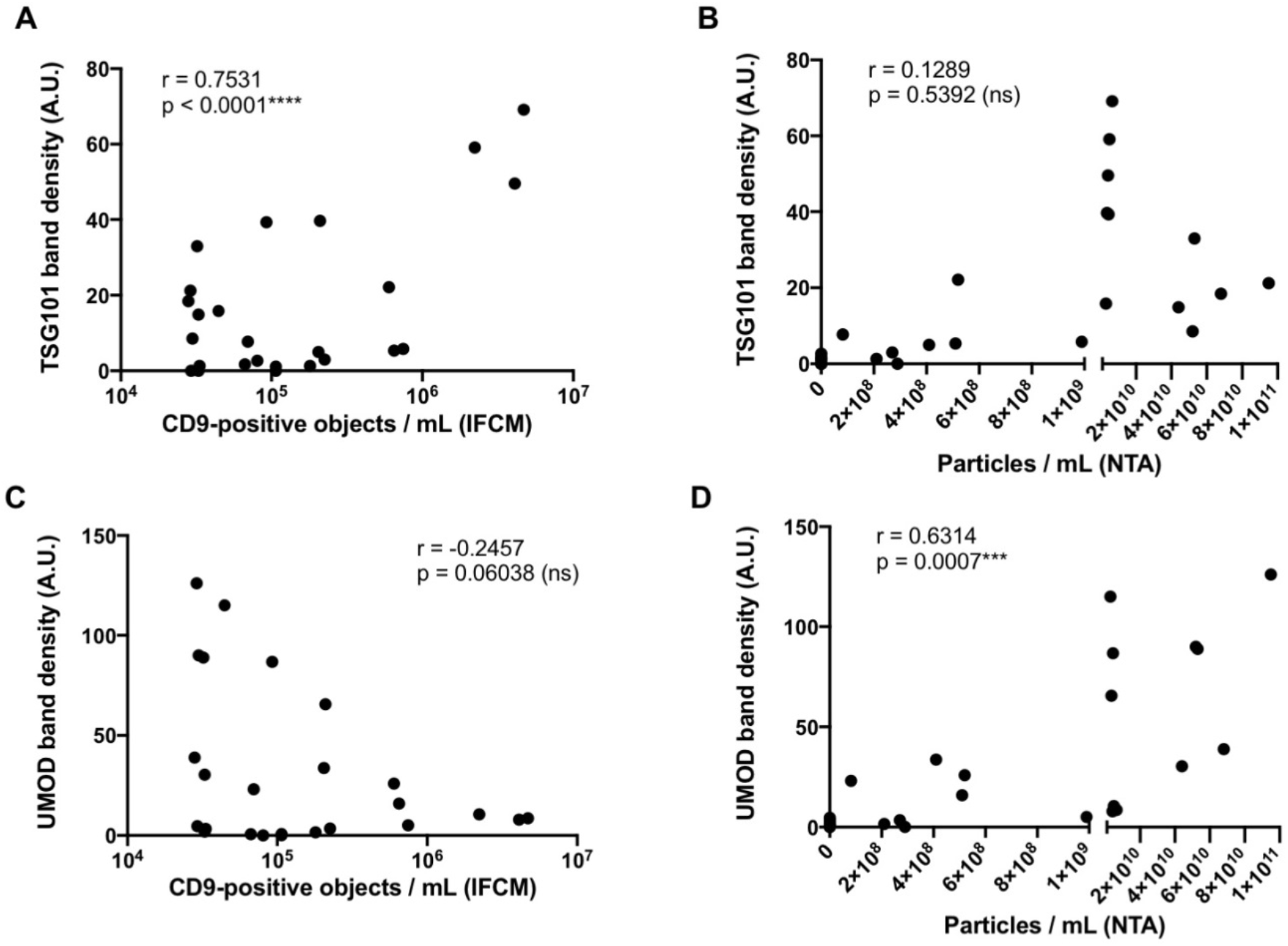
IFCM but not NTA in its traditional form is appropriate for the evaluation of EV preparation methods – IFCM data correlate with results of TSG101 Western blots and NTA data with results of UMOD Western blots. Correlation analyses of acquired Western blot band intensities (TSG101 and UMOD) with CD9^+^ object numbers as determined by IFCM or with average particle numbers as determined by NTA (n = 25) applying Pearson’s correlation coefficient analysis. **A** Correlation analysis of CD9^+^ objects and TSG101 band intensities. **B** Correlation analysis of particle numbers and TSG101 band intensities. **C** Correlation analysis of CD9^+^ objects and UMOD band intensities. **D** Correlation analysis of particle numbers and UMOD band intensities.

### Electron microscopy supports UF-SEC as a suitable method for uEV preparation

To assess the morphology of uEVs that had been prepared with the different methods, TEM analyses were performed (Fig. 7). sEV-sized objects were detected in all preparations. Consistent to the IFCM and WB results, the lowest number of EV-sized objects was found in PEG-UC samples. In samples that were prepared by UC, either with the PEG-UC or the UC-SEC method, detected EV-sized objects were frequently aggregated. Samples that were prepared with the ExoEasy or the UF-SEC method contained higher numbers of non-aggregated EV-sized objects. In UF-SEC samples, these objects showed the EV-typical cup-shaped morphology. Thus, results of the TEM analyses substantiate the result of the IFCM and WB data, indicating UF-SEC apparently allows much better preparation of uEVs compared to all other methods tested here.

**Fig. 7.**
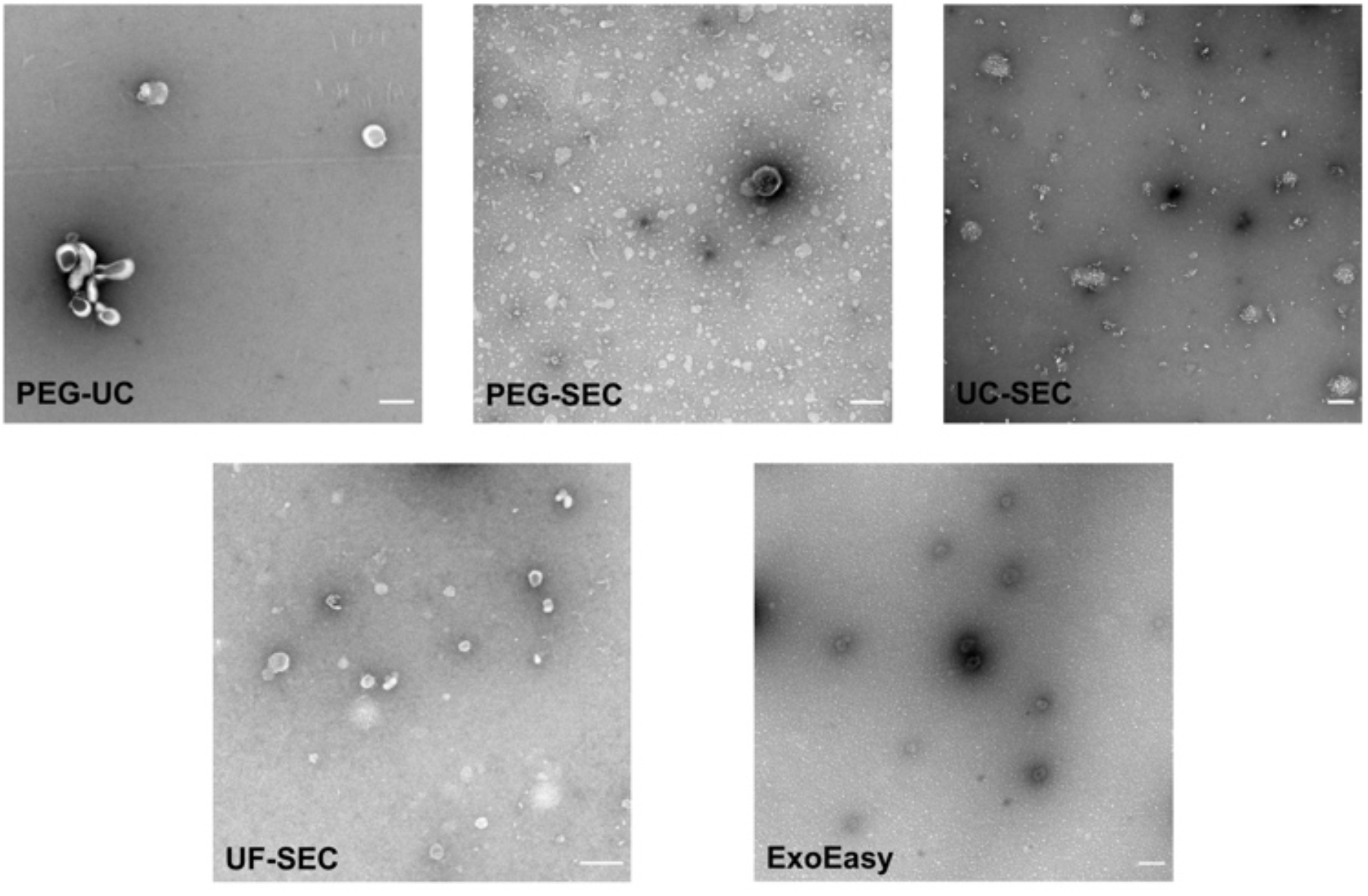
UF-SEC preparation allows recovery of cup shaped uEVs. Representative images of transmission electron microscopic analysis of uEV samples obtained with the different methods as indicated. Size analyses were performed with the ImageJ software. Mean diameters of the EV-like objects were calculated between 67.97 nm (UF-SEC) and 94.73 nm (PEG-UC). Scale bar: 0.2 µm.

## Discussion

Urine is an ideal source of biomarkers due to its non-invasive collection. Apparently, uEVs reflect physiological conditions of the kidneys and the urinary tract and have increasingly been reported as a novel class of urine biomarkers for the diagnosis of kidney diseases (4, 36). However, reflecting the situation in the whole EV field, EV enrichment and detection strategies are still not optimal. The efficiency of EV preparation protocols and the reliability of EV analysis devices remain under discussion, certainly aggravating translation of EVs as biomarkers into daily routine. Here, upon applying reported uEV isolation strategies, we have prepared uEVs from void urine of healthy donors and compared obtained samples using different analysis technologies. As an important finding of our study, we report incongruencies of NTA and IFCM data. While the intensities of WB TSG101 bands correlate with the numbers of CD9^+^ objects measured by IFCM, the intensity of the UMOD bands correlated with particle numbers recorded in NTA. By far, the highest recoveries of CD9^+^ objects from more concentrated void urines were found in UF-SEC samples while highest particle numbers were recovered in ExoEasy followed by UC-SEC samples, the latter having been described as feasible uEV preparation method before (15, 19). Supporting the CD9 data, UF-SEC samples also revealed the highest contrast between the TSG101 and UMOD bands in Western Blots, independent of whether more concentrated or more aqueous void urine samples were processed. UF-SEC samples also contained the highest number of cup-shaped vesicle-like objects. Thus, we conclude that among the methods tested here, UF-SEC provides the best method for uEV preparation.

UF-SEC has been introduced as a feasible uEV preparation method in 2015 (37). Until now, only very few reports have been published that comprehensively compared the accuracy of UF-SEC to other uEV preparation methods applied in the field. A recent study which was conducted in parallel to our investigations and has just been published compared the efficiency of four EV preparation methods (23). Although the authors used NanoFCM in parallel to classical NTA, they focused their conclusions mainly on particle numbers and purities assessed as particle per protein ratio. Upon comparing UF-SEC, UC, polymer-based precipitation with a commercial reagent and Exodisc microfluidics for the preparation of uEVs, the authors identified Exodisc as the best method followed by UF-SEC (23). However, according to WB, the highest EV marker intensities were apparently recovered in the UF-SEC sample. In good agreement with our findings, much less EV marker proteins were recovered in the UC sample and no EV marker protein in the precipitation sample. In contrast to the procedure we have used for IFCM analyses, for the NanoFCM analyses, the authors had to remove unbound antibodies by a subsequent washing and UC-based precipitation step and thus did not discuss the data quantitatively. Since we indeed observed significant EV loss during UC (27), we share their view that quantitative analysis following a UC-based washing procedure needs to be performed with care. However, combined with the results of our study, we would carefully like to conclude that also in their hands UF-SEC is the best method being used, slightly better than the Exodisc device but far better than UC and polymer precipitation. Notably, with the NanoFCM, in good agreement to our IFCM data, the authors detected much more CD9^+^ than CD63^+^ and CD81^+^ objects in their uEV samples (23), highlighting CD9 as an important uEV marker protein.

Based on small particle recovery, a previous study recovered most small particles (50-150 nm) in processed void urine samples following UC-SEC (19). Slightly more small particles were recovered following UC-SEC than UF-SEC samples. Consistent to our study and that of Dong and colleagues (23), PEG and PEG-SEC resulted in low small particle yields (19). In terms of purity, UC samples contained much more protein than UC-SEC and UF-SEC samples, the latter two being in a comparable range. Since UC-SEC samples contained more small particles than UF-SEC the authors judged UC-SEC as the best of their tested methods (19). Despite the information that exosomal marker protein contents were below the detection levels in PEG and PEG-SEC samples but detectable in the other samples, WB data of the method comparison are not presented.

In agreement with our results, polymer or PEG precipitation, which we qualified as a very reproducible method for EV preparation from animal sera or conditioned cell culture media (30), is obviously not appropriate for the preparation of urinary EVs (19, 23). Efficacies of PEG precipitation procedures depend on several parameters including pH, salt and protein concentrations (38), which are apparently not in the permissive range in void urine samples.

The provided and discussed data demonstrate that the interpretation of method comparisons largely depends on the selection of the analysis strategies and the weighting of the results obtained with each of the selected techniques. Underestimating the presence of small non-vesicular particles in classical EV preparations, we and others have introduced NTA as an “exosome” characterization and quantification device (24, 25) which was quickly adopted by the field. Until today, particle quantification by NTA or other particle quantification devices is considered as an essential part of experiments fulfilling the minimal information for studies of extracellular vesicles (MISEV) criteria (7). However, although the problem of non-vesicular particle contamination has been discussed in several manuscripts (39), many of us may have wrongly interpreted obtained results and selected non-optimal methods. The side-by-side comparison performed here, clearly demonstrates the limitations of traditional NTA. Upon learning that IFCM data for CD9^+^ objects correlate very well with TSG101 WB band intensities and NTA data rather with the intensity of UMOD bands, it becomes obvious that NTA is not an EV-specific analysis device but – as expressed by its name – quantifies particles in liquid samples. Since NTA data correlate significantly with the UMOD content in obtained samples, our data imply that at least a proportion of co-prepared UMOD is measured as particles with comparable size ranges than sEVs as it has already been reported for IgG immunoglobulins, myosin aggregates and alpha-synuclein (40, 41). In a related manner, McNicholas and colleagues have previously reported that NTA analyses of uEV preparations of macroalbuminuric disease patients are severely confounded by the presence of albumin. Furthermore, like in our study, their NTA and WB data were incongruent to each other (29).

Questioning the reliability of NTA in its traditional form, care should also be taken when storage conditions for EV-containing samples, including biofluids, are explored. Given the ability of IFCM to detect stained uEVs in unprocessed samples, we observed severe impacts of the storage condition of preprocessed void urine samples on the CD9^+^ uEV recovery after storage with the lowest loss when samples were stored at -80°C and the highest loss – more than 90% – when they were stored at -20°C. Notably, these findings correspond well to WB results of a previous study, which amongst others had investigated impacts of the void urine storage at different temperatures (42). In contrast, NTA analyses of void urine samples failed to detect any storage temperature-dependent particle losses (43). Altogether, the data demonstrate that NTA in its traditional form cannot discriminate between EVs and non-EV particles.

Thus, as confirmed by our data, methods with the highest EV recovery may not provide samples with the highest particle concentration. Accordingly, we would like to recommend the re-evaluation of former method comparisons, especially when conclusions were mainly based on particle recoveries. Like in this study, other methods than those allowing highest particle recoveries may need to be selected as the methods of choice for more efficient EV preparations. This discrepancy demonstrates the urgent need for next generation EV analysis technologies and devices. Indeed, advanced NTA devices have been developed which allow specific tracking of fluorescence labelled particles. Tracking results, however, depend on the labelling efficacies of fluorescent dyes or fluorochrome-labeled antibodies, both providing their own challenges (33). In addition to advanced NTA and IFCM, flow cytometers have been developed to efficiently record single objects in size ranges of sEVs such as the NanoFCM device (44). Based on plasmon resonance, the NanoView device can also analyze the presence of sEVs at the single object level (45). Moreover, a novel direct stochastic optical reconstruction (dSTORM) device, the Nanoimager, allows single sEV analyses (46). In summary, all of these second-generation devices or techniques have their own strengths and weaknesses, but certainly will help to optimize EV preparation protocols and increase our overall understanding about EVs in various body liquids and cell culture supernatants.

## Conclusion

The EV field is progressing almost exponentially and provides plenty of novel diagnostic and therapeutic opportunities. Furthermore, EV studies will certainly help to increase the basic understanding of physiological and pathophysiological processes. Despite the increasing amount of knowledge in the recent decade, however, the methods for EV preparations and analyses remain to be optimized. As demonstrated here, the results generated by second generation analysis devices may question interpretations of past findings and may help to identify experimental weaknesses in current EV preparation technologies. Upon recognizing existing pitfalls and limitations, new technical challenges will arise and help to evolve the field more accurately. Here, we have used IFCM to identify UF-SEC as a powerful method to enrich uEVs from void urine.

## Geolocation Information

Essen, Germany.

## Acknowledgements

The authors would like to thank Dr. Uta Erdbrügger and Dr. Luca Musante (University of Virginia, Charlottesville, VA, USA) for the constructive discussion and criticism of the manuscript.

