## Supplementary Material for "Single extracellular vesicle analysis performed by imaging flow cytometry in contrast to NTA rigorously assesses the accuracy of urinary extracellular vesicle preparation techniques"

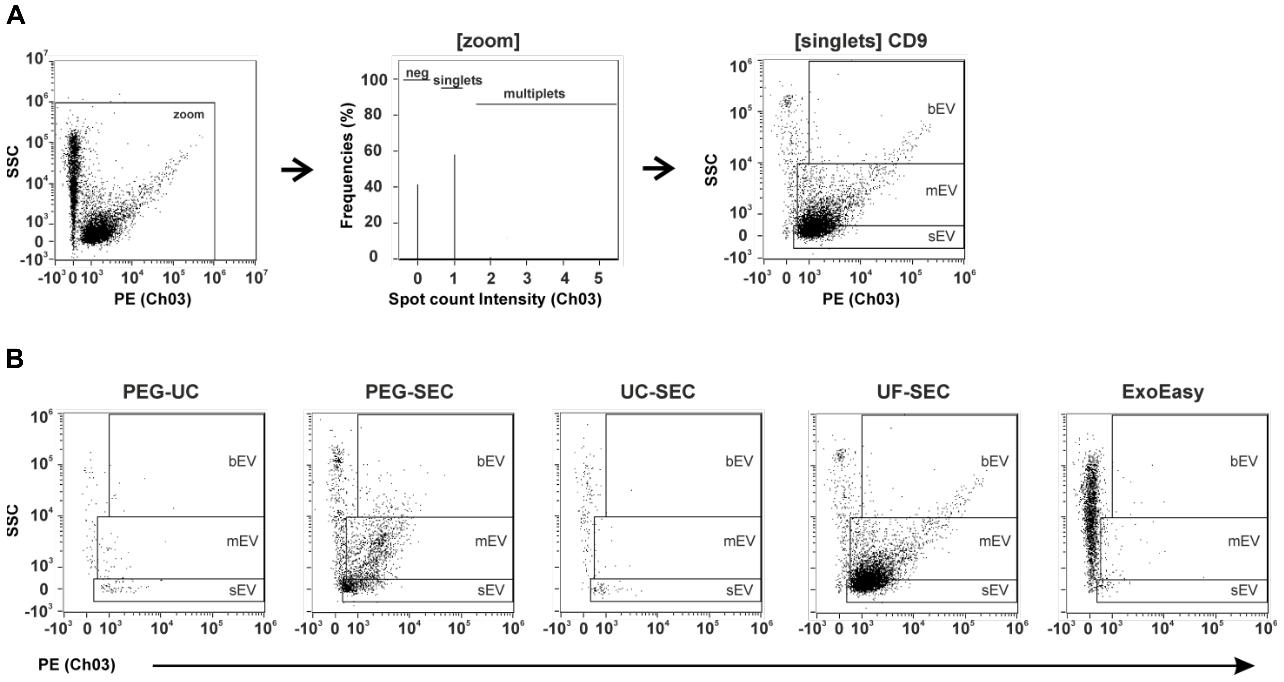


**Supplementary Figure S1. A** Gating strategy for the detection of CD9$^{+}$ objects using IFCM. At first all recorded objects were plotted regarding their side scatter signals (SSC) and their fluorescent intensities following anti-CD9 antibody staining (1$^{st}$ plot). For downstream analyses, coincidences (swarm detection) and objects lacking any fluorescent signal were discriminated from single fluorescent objects (singlets). Only singlets were considered in all downstream analyses. Singlets were also plotted in SSC to fluorescent intensity diagrams. Based on the SSC intensity, three different object subgroups were discriminated: small EVs (sEV), medium-sized EVs (mEV) and big EVs (bEV). **B** Representative flow plots of uEV samples prepared with all five methods.

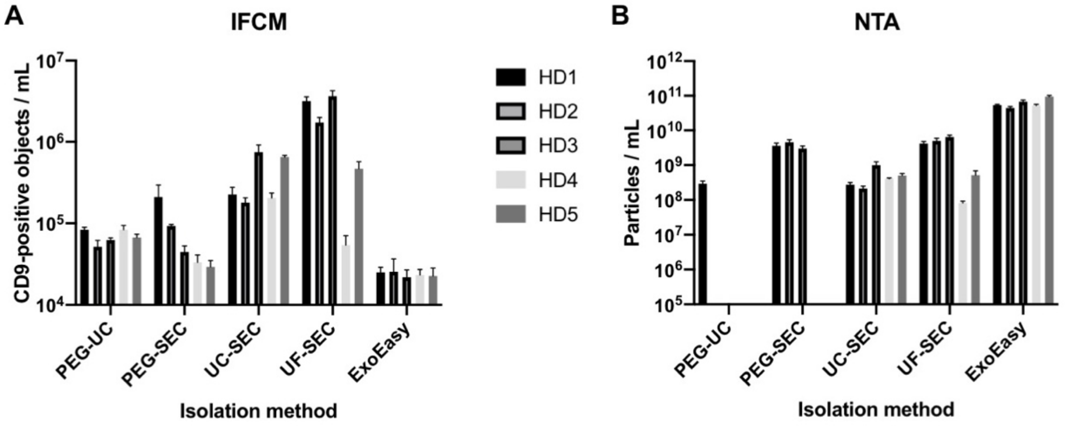


**Supplementary Figure S2.** Detail of Fig. 4: IFCM and NTA results of all prepared uEV samples measured by IFCM (**A**) or NTA (**B**), respectively. Notably, HD1-3 corresponding to the data of the uEV samples of the more concentrated void urine samples and HD4-5 to those of the more aqueous void urine samples. Error bars in **A** represent the variance of three independent IFCM analyses of each sample and those in B the variance of the data measured at 11 positions in NTA.

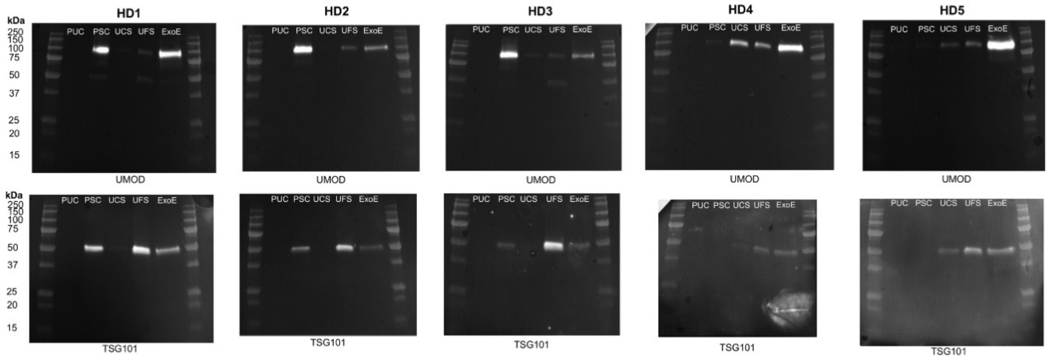


**Supplementary Figure S3. Images of the TSG101 and UMOD Western blots of the uEV preparations obtained from all void urine samples with the different applied methods.** Sample loading of -80°C stored uEV samples was adjusted to volume equivalents of the initial void urine sample. Following sample separation under reducing conditions membranes were sequentially probed with anti-TSG101 and anti-UMOD antibodies (without stripping). HD = healthy donor sample, PUC: PEG-UC; PSC: PEG-SEC; UCS: UC-SEC; UFS: UF-SEC; ExoE: ExoEasy. HD1-HD3 Western blots depict the uEV samples obtained from normal void urine samples and HD4 and HD5 that of the aqueous urine samples.

| Speed | Centrifuge | Manufacturer | Rotor Name | Rotor Type |
|---|---|---|---|---|
| ≤ 4,000 x *g* | Heraeus Megafuge 16R | ThermoFisher Scientific Osterode, Germany | TX-400 | SW |
| 10,000 x *g*<br>17,000 x *g* | Sorvall RC6+ | ThermoFisher Scientific Osterode, Germany | HB-6 | SW |
| > 100,000 x *g* | Optima XPN-80 | Beckman Coulter Krefeld, Germany | Type 50.4 Ti | FA |

**Supplementary Table S4.** Data about the centrifuges, rotors and the centrifugation speed. SW = swing-out rotor, FA = fixed-angle rotor.

| Antibody | Clone, cat. # | Application | Manufacturer | Dilution |
|---|---|---|---|---|
| CD9-PE (mouse) | MEM-61, 1P-208-T100 | IFCM | Exbio, Vestec, Czech Republic | 1:100 |
| CD9 (mouse) | VJ1/20 | WB | F. Sánchez-Madrid* | 1:1,000 |
| CD63-APC (mouse) | MEM-259, 1A-343-T100 | IFCM | Exbio, Vestec, Czech Republic | 1:100 |
| CD81-FITC (mouse) | JS-64, B25329 | IFCM | Beckman Coulter, Indianapolis, IN, USA | 1:100 |
| TSG101 (rabbit) | Polyclonal HPA006161 | WB | Atlas Antibodies, Bromma, Sweden | 1:1,000 |
| THP (UMOD) | B-2 sc-271022 | WB | Santa Cruz, Dallas, TX, USA | 1:200 |
| Goat anti-rabbit IgG-HRP | Polyclonal sc-2004 | WB | Santa Cruz, Dallas, TX, USA | 1:10,000 |
| Rabbit anti-mouse IgG-HRP | Polyclonal sc-358914 | WB | Santa Cruz, Dallas, TX, USA | 1:10,000 |
| Mouse IgG2a-FITC | S43.10 130-113-833 | IFCM | Miltenyi Biotec, Bergisch Gladbach, Germany | 1:250 |
| Mouse IgG1-PE | MOPC-21 555749 | IFCM | BD Biosciences, Heidelberg, Germany | 1:250 |
| Mouse IgG1-APC | MOPC-21 400122 | IFCM | BioLegend, San Diego, CA, USA | 1:250 |

**Supplementary Table S5.** List of the applied antibodies and isotype controls including information about their clone and ordering number as well as the applied dilution factor. *The CD9 antibody (clone VJ1/20) was kindly provided by Dr. F. Sánchez-Madrid, Madrid, Spain.

| Laser [nm] | used Power [mW] | max. Power [mW] | Filter [nm] |
|---|---|---|---|
| 375 | 70 | 70 | - |
| 488 | 100 | 100 | FITC (Ch02) 480-560 |
| 561 | 200 | 200 | PE (Ch03) 560-595 |
| 648 | 150 | 150 | APC (Ch11) 642-745 |
| 785 (SSC) | 70 | 70 | SSC (Ch06) 756-780 |

**Supplementary Table S6.** Applied laser settings for imaging flow cytometry.

| | Ch1 | Ch2 | Ch3 | Ch4 | Ch5 | Ch6 | Ch7 | Ch8 | Ch9 | Ch10 | Ch11 | Ch12 |
|---|---|---|---|---|---|---|---|---|---|---|---|---|
| Ch1 | 1 | 0.029 | 0.042 | 0 | 0 | 0 | 0 | 0 | 0 | 0 | 0.002 | 0 |
| Ch2 | 0.051 | 1 | 0.05 | 0 | 0 | 0 | 0 | 0 | 0 | 0 | 0.002 | 0 |
| Ch3 | 0 | 0.13 | 1 | 0 | 0 | 0 | 0 | 0 | 0.02 | 0 | 0.002 | 0 |
| Ch4 | 0 | 0.064 | 0.49 | 1 | 0 | 0 | 0 | 0 | 0 | 0 | 0.003 | 0 |
| Ch5 | 0 | 0.017 | 0.155 | 0 | 1 | 0 | 0 | 0 | 0 | 0 | 0.074 | 0 |
| Ch6 | 0.015 | 0.02 | 0.04 | 0 | 0 | 1 | 0 | 0 | 0 | 0 | 0.01 | 0 |
| Ch7 | 0.023 | 0.003 | 0.003 | 0 | 0 | 0 | 1 | 0 | 0.015 | 0 | 0.024 | 0 |
| Ch8 | 0 | 0.032 | 0.008 | 0 | 0 | 0 | 0 | 1 | 0.012 | 0 | 0.023 | 0 |
| Ch9 | 0 | 0.004 | 0.084 | 0 | 0 | 0 | 0 | 0 | 11 | 0 | 0.024 | 0 |
| Ch10 | 0 | 0.002 | 0.041 | 0 | 0 | 0 | 0 | 0 | 0.084 | 1 | 0.028 | 0 |
| Ch11 | 0 | 0.001 | 0.012 | 0 | 0 | 0 | 0 | 0 | 0.025 | 0 | 1 | 0 |
| Ch12 | 0 | 0 | 0.003 | 0 | 0 | 0 | 0 | 0 | 0.013 | 0 | 0.125 | 1 |

**Supplementary Table S7.** Applied compensation matrix for imaging flow cytometry.
